# Say what I mean – Expectancy effects in the EEG during joint and spontaneous word-by-word sentence production

**DOI:** 10.1101/782581

**Authors:** Tatiana Goregliad Fjaellingsdal, Diana Schwenke, Stefan Scherbaum, Anna K. Kuhlen, Sara Bögels, Joost Meekes, Martin G. Bleichner

## Abstract

Our aim in the present study is to measure neural correlates during spontaneous interactive sentence production. We present a novel approach using the word-by-word technique from improvisational theatre, in which two speakers jointly produce one sentence. This paradigm allows the assessment of behavioural aspects, such as turn-times, and electrophysiological responses, such as event-related-potentials (ERPs). Twenty-five participants constructed a cued but spontaneous four-word German sentence together with a confederate, taking turns for each word of the sentence. In 30% of the trials, an unexpected gender-marked article was uttered by the confederate. To complete the sentence in a meaningful way, the participant had to detect the violation, (possibly) inhibit a prepared response, and retrieve and utter a new fitting response. We found significant increases in response times after unexpected words and – despite allowing unscripted language production and naturally varying speech material – successfully detected significant N400 and P600 ERP effects for the unexpected word. The N400 EEG activity further significantly predicted the response time of the subsequent turn. Our results show that combining behavioural and neuroscientific measures of verbal interactions while retaining sufficient experimental control is possible, and that this combination provides promising insights into the mechanisms of spontaneous spoken dialogue.

## Introduction

The exchange between two persons conversing is stunningly fast: interlocutors take turns speaking and listening at a rapid rate, requiring them to produce and process language simultaneously. The time needed for producing an utterance is commonly longer than the response times observed during natural conversations ^1^. This speed can be accomplished by forming expectations, for example of the length of a turn and the meaning of the utterance ^2,3^, and then preparing the own response based on these expectations ^4^. When expectations are not met, however, language processing is often slowed down ^5,6^. Aside from the time needed to process the unexpected event, interlocutors may need additional time to adapt a prepared response to the new unexpected context.

To understand the effect of expectations and expectation violations during interactions, not only behavioural but also neural underpinnings might provide a valuable frame. There is, indeed, a major interest in moving towards a neuroscience of social interaction ^7–10^. Interactions, however, are characterized by their openness, while neuroscientific devices impose major constraints to measure sensible brain data. It is challenging to bring these two together ^11^.

One setting in which interactive patterns can be observed in an experimentally controllable environment is the word-by-word exercise from improvisational theatre ^12^. In this game, two persons construct a story together by taking turns for each word. A high degree of coordination is necessary, as the interacting partners have to adapt to each other turn-by-turn in order to produce a meaningful sentence. We believe this level of coordination is achieved by forming expectations about the partner’s next utterance. Similar to natural interactions, one can observe that when expectations are not met the player hesitates to produce the next turn. The word-by-word setting further keeps the principal turn-taking structure of natural interactions intact while giving the possibility to manipulate systematically whether the preceding turn prompts an unexpected sentence completion.

Electroencephalography (EEG) has a high temporal resolution, which can capture the fine temporal structure of verbal interactions. The clear structure of the word-by-word paradigm lends itself for EEG recordings as it ensures valid segmenting of the time periods of interest. EEG studies including semantic expectancy violations have mainly reported modulations of the N400 event-related-potential (ERP), linked to semantic processing, and the P600 ERP linked to syntactic processing ^5,13–15^. The N400 effect is characterized by an amplitude modulation in the averaged EEG approximately 400 ms after an unexpected compared to an expected word, indexing sensitivity to semantic expectancy ^16–23^. The N400 is often followed by a so-called late positive complex or P600 that has been associated not only with syntactic analysis but also overall re-analysis ^15,24^ and even semantics ^25,26^.

In the present word-by-word study, we want to allow unscripted – though controlled – language production of the participant while producing a sentence interactively with a confederate. We measure the underlying electrophysiological activity while systematically manipulating whether the participants’ expectations are met or not. For this purpose, we make use of pictures showing objects that have more than one naming option, i.e., synonyms, with different gender-marked articles in German. To control utterances of participants without making them read aloud, we cued each sentence with a written verb and a picture of an object, both of which had to be included in the sentence. The confederate inserted unexpected sentence continuations (i.e., articles of unexpected gender), where the participant not only had to process the unexpected event but also needed to retrieve and produce a new response deviating from his or her preferred object name.

Behavioural studies on speech production suggest that final word selection among lexical competitors takes place rather late, around 300 ms after picture onset, meaning that multiple candidates are activated at first ^27–29^. Neurophysiological studies of overt language production have found that this lexical access manifests as a positive deflection around 200 ms after event onset ^30–33^. During speech comprehension, discourse (i.e., sentential context) can aid in pre-activating appropriate lexical representations, leading to behavioural costs and differing neurophysiological responses when expectations are violated ^5^. Accordingly, we hypothesize that during interactive sentence production, such as in the word-by-word paradigm, a specific lexical candidate (e.g., the preferred naming of the object on the picture) will be pre-activated to produce the next turn as fast as possible ^1^. Based on this pre-activation, the player will make predictions about the co-player’s preceding turn. When the predictions are violated (i.e., when the confederate utters an unfitting gender-marked article with respect to the pre-activated noun), the player will need to recover by activating one of the less preferred lexical alternatives in order to produce a meaningful sentence.

The successful completion of the word-by-word task entails language comprehension, language production, and for instances of expectancy violations their detection along with the inhibition of a possibly pre-activated response. Our objectives here can be summarized as: (1) testing the feasibility of measuring sensible neural correlates along with behavioural markers in an interactive setup that allows unscripted language production, and (2) measuring some of the neural processes related to successful interactive language use and repair during spontaneous sentence production. For unexpected continuations (i.e., a different gender-marked article from the preferred object noun), we predicted an N400 and P600 ERP effect. Further, we predicted increased turn-times for the next response after encountering an unexpected article. To our knowledge, this is the first study to target interactive language use during EEG measurement with a paradigm that allows such a dynamic sentence production.

## Methods

### Participants

Twenty-five healthy right-handed participants took part in this study. All were German-native speakers, students at the University of Oldenburg and were financially compensated for participation. Written informed consent was collected prior to participation. The study was approved by the local ethics committee of the University of Oldenburg. Research was conducted in accordance with the relevant guidelines. One participant was excluded from analysis due to technical difficulties during the recording. The remaining participants were on average 23.6 ± 2.5 years old (14 female).

### Paradigm

The word-by-word paradigm is inspired by improvisational theatre, where two persons act as one to produce a sentence together, taking turns for each word. In the current experiment, participants’ task was to construct a correct four-word sentence – *Subject-Verb-Article-Object* – taking turns for each word, together with a confederate, e.g., *‘Tina-sieht-das-Sofa (Tina sees the sofa)’*. The participants were cued with a written word (the verb of the sentence) and a picture showing the object of the sentence. The confederate was cued with the whole sentence written out. The participant always uttered the verb and object of the sentence (i.e., second and fourth word). The confederate always uttered the subject and article of the sentence (i.e., first and third word). The paradigm relies on the fact that the German language has three grammatical genders for nouns with accompanying gender-marked definite articles (neuter *‘das’*, feminine *‘die’*, masculine *‘der’*). Hence, we could manipulate whether the confederate’s use of the article matched the participant’s individual preferred naming response (see next paragraph) for the sentence-final object. In 30% of the experimental trials, the confederate uttered an unexpected article (e.g., *‘die’* from *‘die Couch’*) which did not match the participant’s preferred object naming (e.g., *‘das’* from *‘das Sofa’*). In order to construct a correct sentence, the participant had to adjust to the unexpected article and produce the synonym that fitted this article (i.e., *‘Couch’* instead of the preferred naming *‘Sofa’*; see Fig. 1 C).

**Figure 1:**
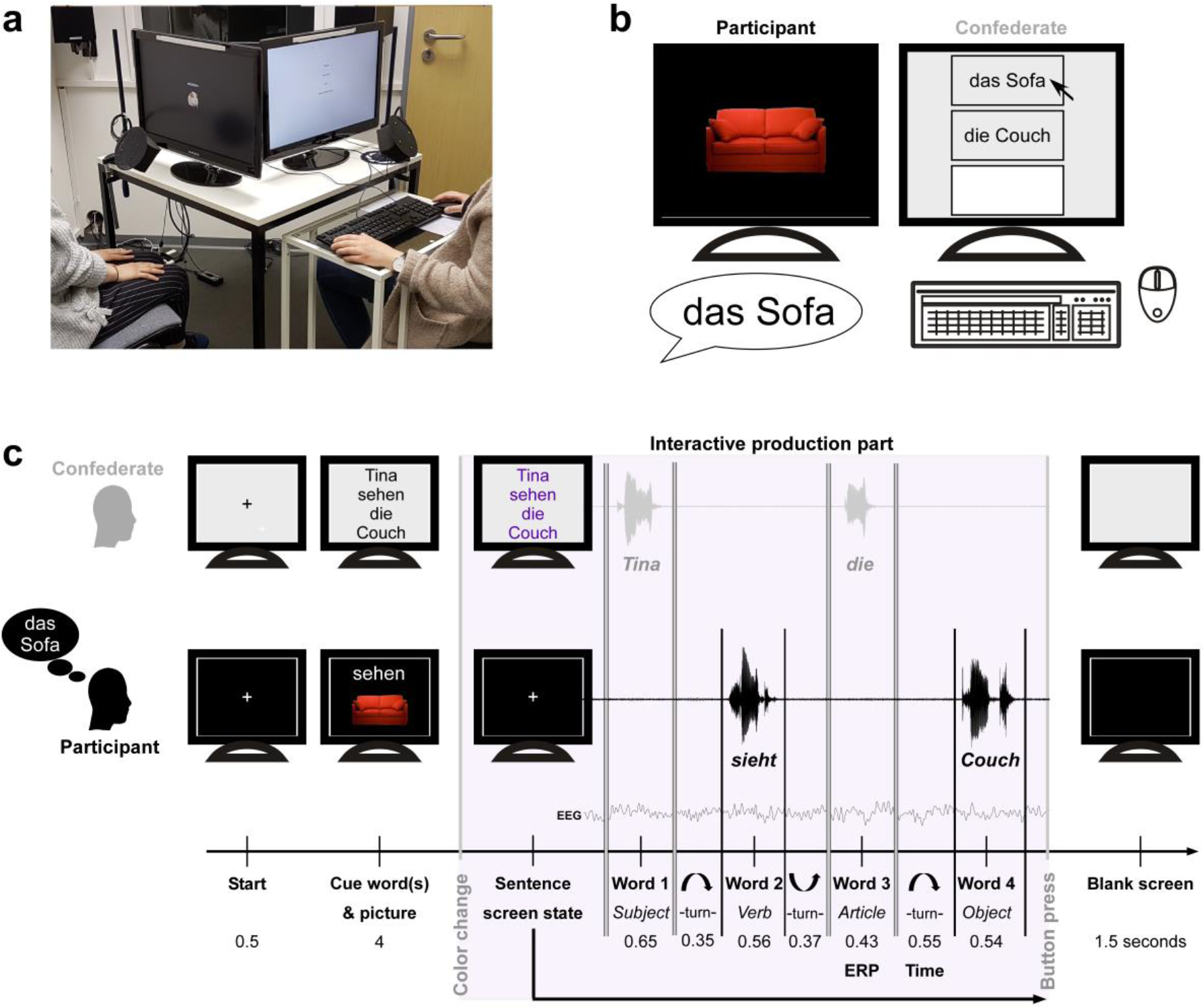
**(a) Setup**: Participant with EEG cap sat on the left hand side in front of a computer screen (no study participant is shown). A microphone was placed near her mouth. The confederate sat next to her in front of a second computer screen and a second microphone, as well as a table with keyboard and mouse. **(b) Picture-naming task**: As soon as the participant named the presented picture, the confederate clicked on the respective synonym or typed in a third option. Namings were automatically saved for the experiment. **(c) Word-by-word timeline**: Screen states and streams of participant and confederate. Example of an unexpected trial: participant named the picture *‘das Sofa’*, expects the article *‘das’* and hears the article *‘die’*. Steady presentation times of fixation cross and words and picture are given in seconds. During the interactive production part (lilac box), the participant saw a steady white fixation cross. The confederate started the sentence with word 1 (sentence subject), the participant then uttered word 2 (sentence verb), the confederate continued with word 3 (sentence article) and the participant ended the sentence with word 4 (sentence object). The interactive production part ended with the button press of the confederate. A blank screen was presented between trials. Average word duration for each word of a sentence, as well as turn-times from word to word are given in seconds. The ERP is computed for the expected/unexpected article (word 3) and the response time until onset of word 4 after expected/unexpected article is contrasted. Sentence translation: *“Tina sees the couch.”*

Participants’ preferred object naming was assessed prior to the main experiment by a picture-naming task (see Fig. 1 B) with three non-sequential naming instances. The last (third) naming instance was used as preferred naming within the experiment. Having three naming instances, gave an intra-individual measure of frequency (i.e., how stable the naming within participant was). Comparing participants’ naming instances over the experiment gave an inter-individual frequency measure of picture naming (i.e., how frequent the used naming was over all participants).

### Material

The 144 pictures used in this study are partly taken from Bögels, Barr, and colleagues ^34^. The 94 critical pictures (49 expected, 45 unexpected) have at least two German naming options, i.e., synonym 1 and synonym 2, where each synonym has a different grammatical gender and therefore different German definite articles (e.g., *‘das Sofa’* vs. *‘die Couch’*; see supplementary figure 1). The remaining 50 pictures were selected to have one common German naming option and were used as filler trials.

The written word stimuli consisted of 144 first names (Number of letters: *M* = 5.48 ± 1.48, range 3 to 9; 50% female) and 144 German verbs (Number of letters: *M* = 7.79 ± 1.74, range 5 to 13; non-reflexive). Verbs were selected avoiding common co-occurrences of the verb with a particular object naming.

### Procedure

Participants were screened (to exclude neurological, psychiatric and/or language disorders) via a telephone interview prior to invitation. Candidates who knew the confederate (the same person throughout the study) were excluded. On the day of measurement, the participant first signed informed written consent and filled in two questionnaires (Edinburgh Handedness Inventory ^35^ and a questionnaire on demographic and physiological information). Participants were briefed about the following tasks and that their interacting partner was a confederate.

For the measurement, participant and confederate sat next to each other, each in front of a computer screen (see Fig. 1 A). The confederate additionally had a small desk with a keyboard and mouse. Two microphones (ETM-006, Lavalier Microphone) with an audio pop shield were located on an extension on the table and each one was placed approximately 15-20 cm in front of participant and confederate respectively. To keep the microphone recording as clean as possible, participants were instructed to utter only the words of the experiment and to avoid filling utterances (e.g., ‘ehm’, ‘eh’) and other vocal noises (e.g., laughs, throat clearing). The picture-naming task was programmed in Matlab R2017b. The word-by-word experiment was programmed with the psychophysics toolbox ^36,37^ in Matlab R2017b.

First, a picture-naming task with two naming instances per picture was conducted. Participants were asked to name each picture with its respective definite article. Three practice trials were used to familiarize the participant with the task. The 144 pictures (cropped image on black screen) were shown once in a randomized order and then again in a different randomized order (i.e., 288 trials in total) to the participant. The confederate sat alongside the participant and saved the respective responses by clicking on the predefined synonyms or typing and saving the new naming options (see Fig. 1 B). The participant was not able to see the screen of the confederate (see seating in Fig. 1 A) and was not informed about the task of the confederate.

Thereafter, the EEG cap was fitted and impedances controlled. A second picture-naming task with one naming instance per picture (i.e., 144 trials) was conducted (the same cropped images on black background). The confederate again saved the respective namings, which were used from this second run as target words for the expectancy manipulation in the word-by-word experiment.

Subsequently, the word-by-word experiment (see Paradigm and Fig. 1 C) was conducted. Here, the task was to construct a correct four-word sentence, taking turns for each word. For the participant, each trial started with a fixation cross (0.5 sec), followed by the simultaneous presentation of the written verb in infinitive (white letters) in the upper middle centre and the cropped object picture in the lower middle centre of the screen (4 sec). During the interactive production part, the participant’s screen displayed a steady fixation cross and the confederate’s screen displayed the scripted words of the sentence. The confederate uttered the first word, *‘Tina’* (sentence subject), the participant then uttered the second word, which had to be conjugated, *‘sieht’* (*sees*, sentence verb), the confederate then uttered the third word, *‘die’* (*the*, sentence gender-marked article), and the participant uttered the fourth word, *‘Couch’* (sentence object). The trial was terminated with a button press by the confederate. A blank screen was presented for 1.5 seconds between trials. The article uttered by the confederate could either match the preferred naming of the participant (70% of trials – ‘expected’) or fit an alternative naming with different grammatical gender (30% of trials – ‘unexpected’). We emphasized the importance of producing a correct sentence with the confederate using the verb and picture prior to the trial, without revealing that they would encounter unexpected sentence continuations. Three practice trials (without expectation violations) were conducted to clarify the task. Participants were instructed to keep movement minimal and to use the time of the blank screen between trials for necessary movements. Every 12 trials there was a pause and participants could decide when to continue.

After the experiment, the EEG cap was removed and participants were asked to fill in an evaluation questionnaire. Participants were for example asked to rate how natural the interaction seemed, how pleasant it was, and how pleasant the interaction partner was on a five-point scale from ‘not at all’ to ‘very’ (see supplementary – evaluation results). A complete experimental session lasted around 3 to 3.5 hours.

### EEG Recording

Brain electrical activity was measured with a 96-channel EEG system (BrainProducts, Gilching, Germany). Ag/AgCl electrodes were placed equidistantly with a nose tip reference and centro-frontal ground (Easycap, Herrsching, Germany). Impedances were kept below 20 kΩ. Data were digitized with a sampling rate of 500 Hz. EEG stream, marker stream, and audio streams were recorded synchronously using the Lab Recorder from Lab Streaming Layer ^38^.

### Audio Preprocessing

Word onsets and offsets were first roughly estimated to create epochs around each single word. Next, the audio signal of these epochs was high-pass FIR filtered at 35 Hz and down sampled to 1470 Hz. The envelope was computed (filter length 300) and a low-pass Butterworth FIR filter at 730 Hz was applied. Thereafter, the root mean square (RMS) and cepstrum (using the Voicebox toolbox ^39^) were calculated, and the first and second fundamental frequencies were extracted from the cepstrum (low-pass filtered at 600 Hz). To find the real speech onset, we applied the function ‘findchangepts’ (Matlab toolbox signal processing) on the RMS, which gives a series of markers showing changes in the RMS audio signal. The onset marker was set as valid, if changes were apparent in all calculated signals (RMS, envelope, first and second fundamental frequencies). For speech offset detection, the same procedure was applied with a time-reversed audio signal. Each word segment (from onset to offset) was inspected by ear and adjusted, if necessary.

### Behavioural Analysis

The results of the picture-naming task, where each picture had to be named three times, were assessed. When the same name was used for a picture all three times it received an intra-individual frequency rating of 2, when the same name was used twice it received a rating of 1, and when it was named differently each time it received a rating of 0, indicating that the participant did not have a stable and preferred naming of this object. In addition, we assessed the inter-individual frequency naming per picture across participants and naming instances. For the two predefined synonymic namings, we counted how often each synonym (synonym 1 & synonym 2) was chosen by the participants. All cases in which participants chose another option were combined into a third category (option 3). The respective naming percentage was then computed for each picture (see supplementary figure 1 for an overview). Both measures, intra-individual frequency and inter-individual frequency, were used as additional explanatory variables for the turn-time prediction model (see statistical analysis below). The idea here is that individuals might cope better with expectation violations if they do not have a clear naming preference for the object on the picture.

Average speech durations and word length (number of letters) were calculated for all words. For word 4, the response accuracy was assessed. Trials were classified as erroneous and excluded from the behavioural analyses when the verb was forgotten or when the participant did not utter the fourth word or uttered an ungrammatical and/or wrong word (1.07% of all trials; 1.39 % of critical trials, i.e., 0.51% from congruent and 2.38% from incongruent trials).

Turn-times were calculated from word offset of the previous word to word onset. Turn-times over 5 seconds and below 0 seconds were excluded from further analysis. To assess statistical differences of turn-times from word 3 to word 4 between conditions, a Generalized Linear Mixed Model (GLMM; ^40^) was calculated in R ^41^ with the lme4 package ^42^. To account for non-normally distributed reaction time data, the model was fit with a probability distribution of the Gamma family and an inverse link function ^43^. We included the fixed factors expectancy (expected vs. unexpected), intra-individual frequency (intra-individual frequency of naming: zero repetition vs. one repetition vs. two repetitions), and inter-individual frequency across participants (general frequency of naming of the picture with chosen naming; compare section Paradigm). Both latter factors, intra-individual frequency and inter-individual frequency, could have an effect on the time needed to recover from an expectation violation. To account for a possible interaction between inter-individual frequency and expectancy, an interaction term between these two factors was included in the model. A random slope congruency for the random intercept subject was added to model possible inter-individual differences in effect size of expected to unexpected condition. Furthermore, we included the random intercept factor length of word (i.e., the number of letters) in the model.

### EEG Analysis

Preprocessing was performed with EEGLAB ^44^ in Matlab. For artefact attenuation, we applied extended infomax Independent Component Analysis (ICA) on finite impulse response (FIR) filtered data (1 to 40 Hz). To semi-automatically remove artifactual components, we applied the corrmap toolbox ^45^, as in the standardized procedure explained in Stropahl, Bauer, Debener, & Bleichner ^46^. All outer ring electrodes (13 electrodes) were excluded from further analysis, due to increased muscle artefacts.

For ERP analysis, the data was FIR filtered between 0.1 to 30 Hz and re-referenced to average mastoids. The data was epoched from −500 to 1500 ms around word onset as determined by the microphone signal (see Audio Preprocessing) and baseline corrected from −100 to 0 ms. An automatic epoch rejection was applied, where all epochs exceeding three SDs from the mean signal were removed. Further, we applied an automatic artifactual channel detection (EEGLAB function) and interpolated channels exceeding a kurtosis value of 5. Per participant and grand average ERPs were calculated.

Two ERP components for the critical word (CW; word 3) were of main interest: the N400 effect between 250 and 450 ms (compare early N400 effects for short articles in ^14^) and the P600 or late positive complex effect between 500 and 700 ms (see ^24,47^). The mean number of trials entering statistical analysis for word 3 were 43 and 38, for expected and unexpected conditions respectively (filler trials were excluded from analysis). A Linear Mixed Model (LMM) was calculated in R ^41^ with the lme4 package ^42^ separately for each component and each specified region of interest (ROI; see Fig. 3 for locations) along the midline (1 - anterior, 2 - central, and 3- posterior) and along the quadrants (1 - left anterior, 2 - left posterior, 3 - right anterior, and 4 - right posterior). Fixed factor in each model was expectancy (expected vs. unexpected). We included a random intercept for participant and a random slope expectancy within participant. P-values for expectancy were calculated by comparison of the model with and without expectancy factor.

### Brain-behaviour interaction analysis

To test the interaction between brain and behaviour (compare for example ^48^), a GLMM was calculated in R ^41^ with the lme4 package ^42^. Individual turn-times from offset of word 3 to onset of word 4 were the response variable and fixed factors were the respective mean N400 EEG activity between 250 and 450 ms over the specified ROIs (see Fig. 3) and expectancy (expected vs. unexpected). An interaction term between both factors was added to the model. Similar to the GLMM calculated for the behavioural analysis, random factors included word length (intercept) and expectancy (slope) in participant (intercept).

## Results

### Behavioural results

Participants successfully produced a sentence without grammatical errors and with the predefined verb and object in more than 98% of all trials. In critical trials, participants produced on average a correct sentence in 99.49% of expected trials and 97.62% of unexpected trials. Errors were predominantly present in the first of the twelve blocks, indicating a familiarization effect over the experiment and/or problems of the participant understanding the task of constructing a correct sentence. Three participants were reinstructed after the first block to ensure compliance of the task. Erroneous trials (e.g., sentences where the verb was forgotten, an ungrammatical fourth word was uttered or no fourth word was uttered) were excluded from further analysis. On average, the word duration, i.e., the time spent uttering a word, was 559 ± 12 ms for word 2 and 540 ± 30 ms for word 4 (543 ± 38 ms after expected articles and 548 ± 54 ms after unexpected articles).

Participants needed on average 351 ± 59 ms to produce word 2 (turn-time from offset of word 1 to onset of word 2) and 554 ± 193 ms to produce word 4 (turn-time from offset of word 3 to onset of word 4) over all conditions (filler, congruent, incongruent). Split by critical conditions, participants needed on average after an expected article 405 ± 168 ms and after an unexpected article 958 ± 273 ms to produce word 4 (see Fig. 2 a).

**Figure 2:**
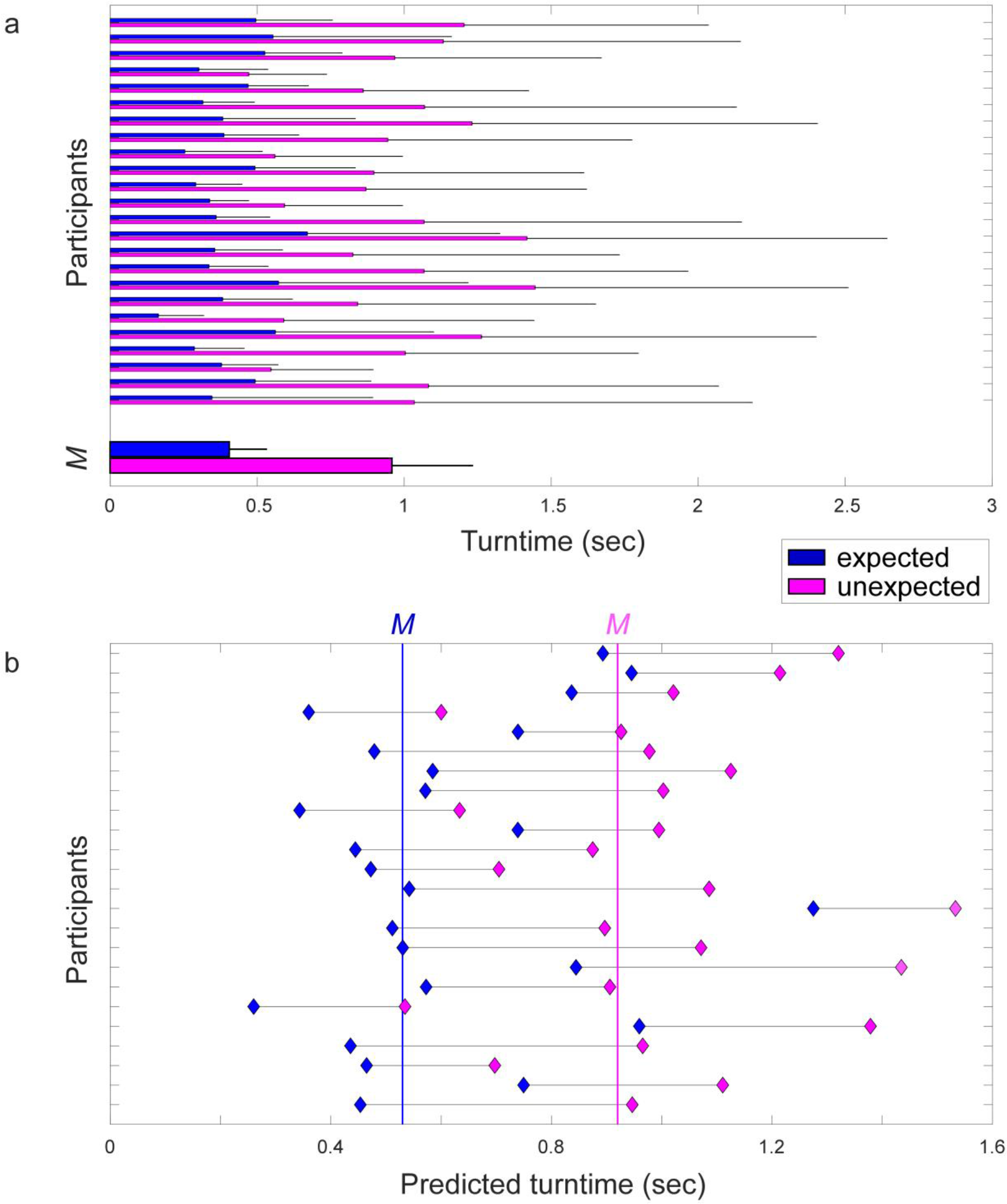
(a) Average turn-times from offset of word 3 to onset of word 4 are shown in seconds for each participant for expected (blue) and unexpected (magenta) conditions plus standard deviation. Grand average turn-time (*M*) is shown below. (b) Predicted average turn-times from offset of word 3 to onset of word 4 from the calculated GLMM, shown for each participant (y-axis) for expected (blue) and unexpected (magenta) conditions. Grand average predicted turn-time for the expected condition is shown as blue vertical line and for unexpected conditions as magenta vertical line.

**Figure 3:**
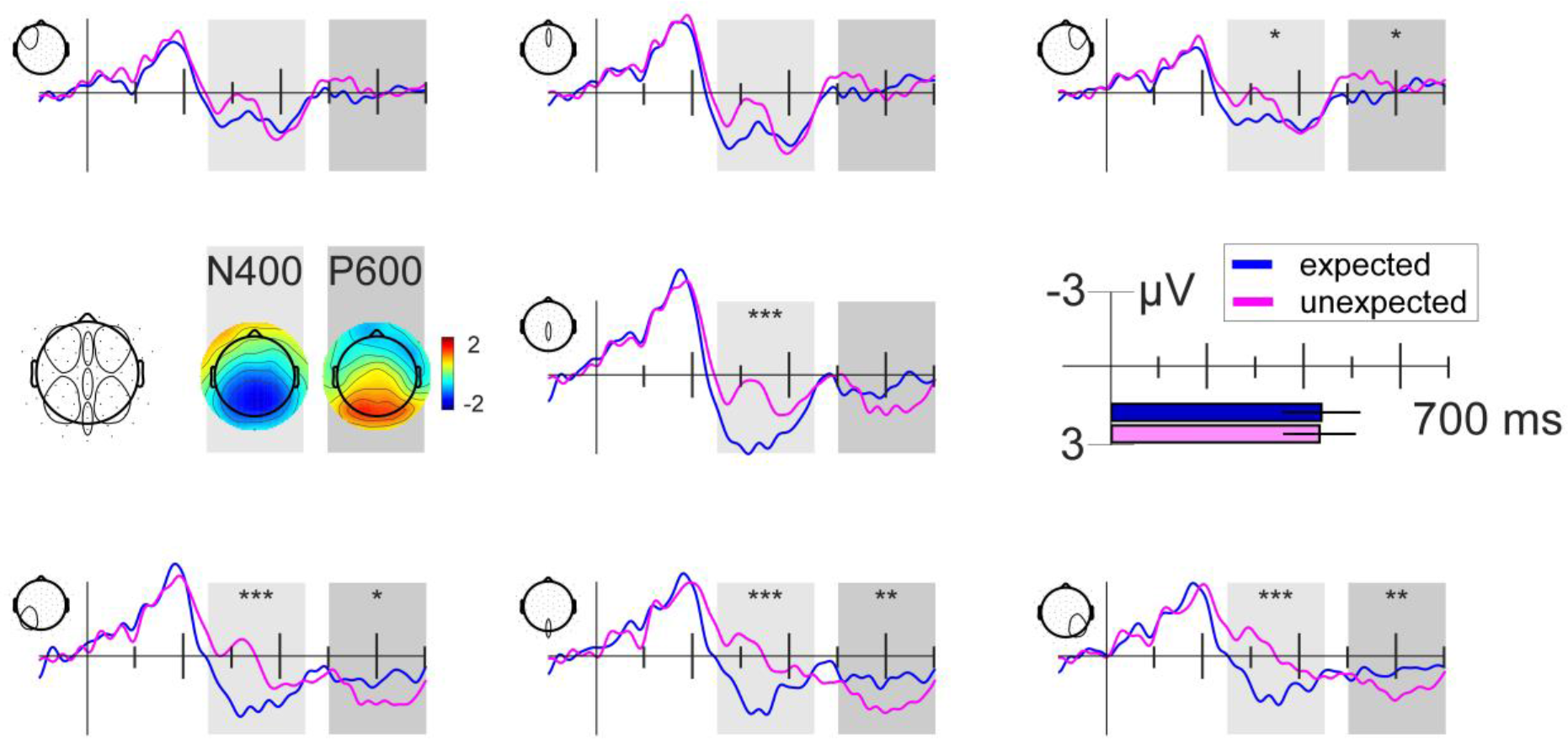
Grand average ERPs of word 3 shown for midline averages and quadrant averages for expected (blue) and unexpected (magenta) conditions (see electrode map for specific regions). Same regions were used for statistical analysis. Difference topographies for the specific time windows are shown. The N400 effect time window is highlighted in light grey (250-450ms). The P600 time window is highlighted in dark grey (500-700ms). Zero point is the onset of word 3, the expected or unexpected article. Mean word duration (of confederate) plus standard deviation is shown below the axis description with the same time scale. * *p* < .05, ** *p* < .01, *** *p* < .001

A GLMM with fixed factors expectancy, intra-individual frequency, inter-individual frequency, and random slope congruency nested in intercept participant, as well as random intercept word length showed that intra-individual frequency of naming (i.e., intra-individual naming for the same picture) had no significant effect on turn-time (*χ*2 (2) = 1.88, *p* = .391). Therefore, it was dropped from the model for better model fit (see supplementary table 1). Results of the final GLMM (see Fig. 2 b) showed that expectancy (expected vs. unexpected) had a significant effect on turn-times (*χ*2 (3) = 83.88, *p* < .001), with increased turn-times for unexpected events (see supplementary table 1). Further, a significant interaction between expectancy and frequency of the preferred naming (i.e., inter-individual naming distribution) was present (*χ*2 (1) = 40.98, *p* < .001), where higher frequencies of a naming lead to decreased turn-times for expected events and increased turn-times for unexpected events. Excluding the nested random slope for congruency in participant significantly decreased model fit (*χ*2 (2) = 62.90, *p* < .001), indicating considerable variation between participants in the effect of the expectancy violation (see Fig. 2 b).

### EEG results

The grand average ERP of word 3 for expected and unexpected conditions shows an N400 effect between 250 to 450 ms with a centro-posterior scalp distribution (see Fig. 3). Further, a P600 is apparent between 500 and 700 ms after word onset with a posterior topography (see Fig. 3).

Statistical analysis confirmed that expectancy had a significant effect on the N400 amplitude for word 3 over five of the seven specified ROIs (see Fig. 3 for electrode locations and see supplementary table 2 for an overview of all results; e.g., posterior midline: *χ*2 (1) = 15.48, *p* < .001, left posterior quadrant: *χ*2 (1) = 35.14, *p* < .001, and right posterior quadrant: *χ*2 (1) = 34.02, *p* < .001). The N400 amplitude was significantly more negative for the unexpected compared to the expected condition. The topographical distribution and statistical result indicate a widespread N400 effect with a posterior maximum (see Fig. 3 and supplementary table 2).

Expectancy also significantly modulated the P600 amplitude for word 3 (see supplementary table 3) over posterior midline (*χ*2 (1) = 8.66, *p* = .003), left posterior quadrant (*χ*2 (1) = 6.40, *p* = .011), right anterior quadrant (*χ*2 (1) = 3.96, *p* = .046), and right posterior quadrant (*χ*2 (1) = 8.34, *p* = .004). The P600 amplitude was significantly more positive for the unexpected condition compared to the expected condition over all mentioned ROIs, except over the right anterior quadrant, where it was more negative for the unexpected condition. These results indicate a posterior topographical distribution of the P600 effect.

### Brain-behaviour interaction results

The GLMM calculated for turn-time from word 3 to word 4, including the mean N400 EEG activity from 250 to 450 ms and expectancy condition as explanatory factors showed that for each specified ROI (anterior, central, and posterior midline, and left anterior, left posterior, right anterior, and right posterior quadrant; see Fig. 3) N400 EEG activity had a significant effect on the resulting turn-time (all *p* < .001, see supplementary table 4). As expected, turn-times were significantly longer in the unexpected than in the expected condition (all *p* < .001, see supplementary table 4). There was no significant interaction between expectancy and N400 EEG activity (all *p* > 0.05, see supplementary table 4).

## Discussion

In this study, two persons jointly produced a sentence, taking turns for each word. The word-by-word paradigm is inspired by a technique used in improvisational theatre, which models various aspects of natural interactions. The paradigm’s structure allows for high experimental control, along with the ability to induce expectation violations during an interaction. These two pillars make the paradigm an effective tool to study neurophysiological (e.g., N400 and P600 ERP) and behavioural effects (e.g., turn-time) during interaction. In the present study, we could successfully induce expectation violations by making a confederate utter a gender-marked article that did not fit the participants preferred object name’s gender that had to be produced in the next turn.

The behavioural findings within this paradigm are what we predicted based on previous research ^4,6,49^ and our own observations from improvisational theatre, as well as everyday experiences of natural interactions. When expectations are not met, the time to produce the next response increased significantly as compared to when the expectations are met. This finding points to the fact that participants pre-activated their preferred object naming in order to produce the next turn fast as possible. However, when they encountered an unexpected (unfitting gender-marked) article, they had to discard the pre-activated lexical entry of their preferred object naming and produce the fitting word. Behaviourally, we can capture the consequences that follow from this repair of a violated expectation. This behavioural effect might still reflect numerous underlying processes, which cannot be disentangled easily.

To further our understanding of the underlying mechanisms during word-by-word interactions, neurophysiological underpinnings can help in disentangling some of the crucial aspects for successful interactions. On the neural level, we predicted two main effects, the N400 and P600, to be modulated significantly by expectancy. This was indeed the case, unexpected articles led to a more negative amplitude of the N400 and a more positive amplitude of the P600. We will discuss in the following paragraphs what their presence in this particular setup can tell us about the interplay of language comprehension and language production during verbal interaction.

The first process observed in the EEG, the N400 effect, is known to index processing of expectation violations in various domains (for an overview see ^22^). Seeing the N400 effect here is consistent with the idea that the participant pre-activates a specific lexical entry and accompanying grammatical gender during the interaction. The N400 effect is the response to encountering an unfitting gender-marked article to this pre-activated entry. Similar to grounding in conceptual pact studies ^50^, i.e., where interacting players agree on a specific term for a specific object, the participant named the pictures pertaining to the objects in the co-constructed sentences prior to the experiment in the presence of the interacting confederate. We deduce that participants ascribed certain expectations to the confederate that she would name the objects the same way they had named them and would therefore utter a fitting gender-marked article. The confederate in fact uttered fitting gender-marked articles to the participants’ expected object names in the majority of the trials (75%), rendering the remaining trials unexpected. Similar N400 effects on the article level (i.e., when the article renders a noun with high cloze probability grammatically incorrect) have been reported in a language comprehension task in 2005 by DeLong and colleagues (see also ^5^). These N400 effects on article level have been interpreted by the scientific community as strong indicators for prediction during language comprehension ^22^, since the article itself constraints the probability of following nouns without defining context in itself. However, DeLong et al.’s findings failed to be replicated in a large-scale replication analysis, suggesting that (phonological forms of) words are not necessarily pre-activated during language comprehension ^51^. Our word-by-word setup combines language comprehension (of the article) with instant language production (of the following noun). We show that in this context a pre-activation of an object form is indeed present, which shows up as an N400 effect on the article when violated. The N400 was even predictive of the resulting turn-time needed to utter the next word. We conclude that pre-activation of a specific lexical entry aids in accomplishing the present word-by-word task in a rapid manner, common to the timely turn-taking structure of natural interactions ^52^. It is an open question, if the pre-activated entry leads to a prepared word (i.e., in the speech production loop) or if it relates to pre-activation that aids in speech preparation after listening to the turn of the partner. In other words, it is unclear if speech production is planned during the turn of the partner *or* after the turn has finished. The later, positive going ERP we observed in the EEG for unexpected conditions could provide information to answer this question.

The classical account of this positivity we see would be that of a P600 ERP that has been linked to syntactic analysis ^24^ and discussed as an index for structural reanalysis, for example regarding semantics ^53^. The P600 or late positive complex often follows an N400 effect (e.g., ^15^). In the present study, the P600 would then reflect the parsing of the unexpected article with a transfer to new retrieval. Sassenhagen and colleagues ^25^ for example found the P600 to be response-aligned to the reaction time of a button press. In this line of argument, the P600 reflects the point, when the event has been fully integrated in the sense-making system, opening the transfer to the most suitable response (be it a button press or speech preparation).

Yet an alternative interpretation of this finding is possible. The late positivity might reflect lexical access: a starting point for the retrieval of the new response after an unexpected article is encountered. In line with speech production research, an early positive EEG component has been discussed as marker for the retrieval of words, which is usually largest at posterior electrode sites ^4,30–32^. It is possible that this lexical retrieval component is present at different time points for the expected vs. unexpected condition (see topographies for the expected condition in supplementary figure 4). For the expected condition, we would then interpret the positivity around 200 ms after the onset of the article as a lexical access response (see supplementary video), indicating speech preparation while still listening to the turn of the confederate (word duration of the article was on average > 400 ms long). In the unexpected condition, lexical access might be disrupted, marked by a larger N400 amplitude, which reflects the processing of the unexpected article. Lexical retrieval is then delayed (or re-activated) at a later stage, for example around 500 to 700 ms or even later, which can overlap with the interpretation of a P600. Given the considerable differences in task demands of earlier EEG studies on speech production and the present study (e.g., picture naming requiring immediate response vs. delayed response) this interpretation is rather speculative. We would encourage future studies to test this interpretation for example by adding a control condition, where participants would listen to the unexpected article without having to produce a response thereafter.

Also in regard of language comprehension, the question remains whether the noun is pre-activated due to the required speech production in the next turn or if the N400 effect on an article can also be found for pure language comprehension scenarios. Assessing the preferred object names of participants prior to a language comprehension task, where they listen to sentences with their preferred and dis-preferred object names could provide a scenario to study this question.

Future studies could moreover target the role of interindividual differences during interactions. We have seen that during the current word-by-word construction participants suffered to a different degree from the expectation violations (see Fig. 2). Such interindividual differences are also visible during joint story building in improvisational theatre. For example, one can observe differences in the response to expectation violations and their repair. Naïve players are often in situations where they cannot come up with a response, while proficient players manage smooth interactions also without knowing the partner beforehand. Grasping these differences with implicit and neurophysiological measures can pave the way to assess the role of learning in coping with unexpected events and its possible transfer to other social situations. The interaction of brain and behaviour in the present study further shows that neural correlates can be predictive of the behavioural outcome.

## Conclusion

Social interactions are complex and marked by multiple levels of processing. Here, we successfully measured neural activity related to linguistic processing during verbal interaction. To our knowledge, this is the first study to measure EEG during expectation violation, where the participant is not only required to comprehend and detect the violation of freely produced speech, but also to inhibit a pre-activated response and retrieve a new response to complete an interactively produced sentence. Our EEG findings revealed two underlying processes of the handling of these expectation violations, with one significant outcome on the behavioural level, i.e., in turn-time. A link of these two measures could be established via a brain-behaviour model, i.e., the N400 effect on the article-level predicted the turn-time to produce the following object noun. We conclude that there is added value in combining both measures, behavioural and neural, to understand the mechanisms of social interactions. This joint assessment was possible with the word-by-word paradigm, which combines verbal interaction with the necessary experimental control.

## Acknowledgments

We thank Marike Maack for her help during data collection. We would also like to thank Reiner Emkes and Florian Wiese for technical support. We are grateful to Prof. Stefan Debener for his support, feedback, and provision of lab facilities.

## Author Contributions Statement

Study conceptualization: TG, MB, SS, DS, AK. Study material and setup: TG, DS, MB, SS, SB. Data analysis: TG, MB, JM. Manuscript writing: TG, MB, SS, AK, SB, JM, DS.

## Data availability statement

Data cannot be made available due to data security requirements of ethic approval.

