## Supplementary information for "Say what I mean – Expectancy effects in the EEG during joint and spontaneous word-by-word sentence production"

### Supplementary Material

#### Inter-individual frequency of naming: synonym 1, synonym 2, and option 3

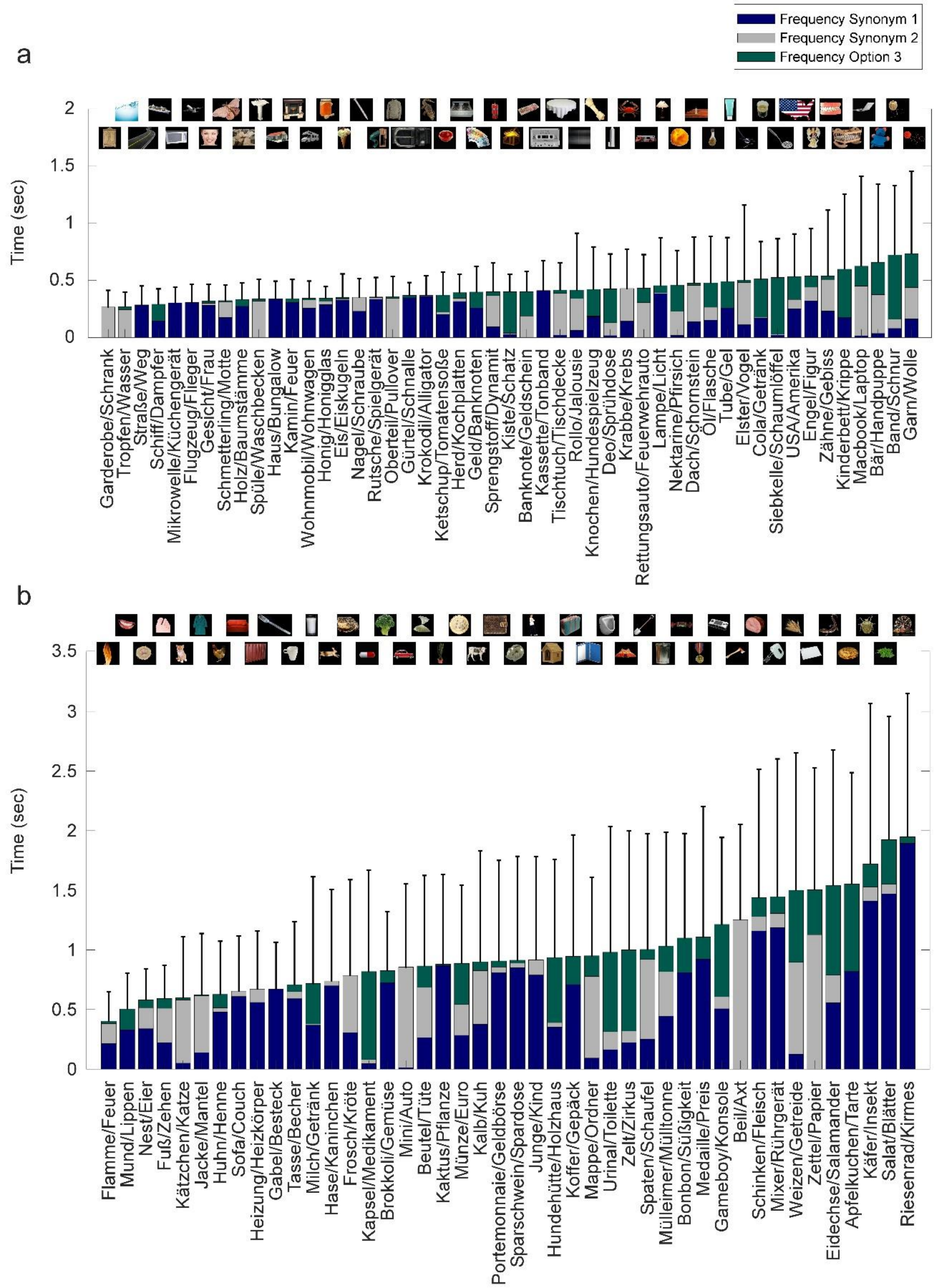


**Supplementary figure 1**: Critical pictures sorted by the grand average turn-time from offset of word 3 to onset of word 4 for the expected condition (a) and unexpected condition (b). Inter-individual frequency of naming is shown percentwise for each picture per bar: Synonym 1 in blue, Synonym 2 in grey, and Option 3 in green. Our predefined (German) synonyms for each picture are shown below each bar (Synonym 1 / Synonym 2). Option 3 was any other naming for the picture, which could also be a composite of the predefined synonyms (e.g., '*Garderobenschrank*' or '*Feuerflamme*').

**Evaluation results**

After the word-by-word experiment, participants filled in an evaluation questionnaire. All participants reported to have understood and followed the instructions of the experiment. Five participants reported to have known the word-by-word game from children or party games prior to the experiment. Eight participants answered the question 'Was there any difficult aspect within the experiment?' saying that it was hard to continue the sentences after articles that did not match their object naming. When asked about the goal of the study (participants only knew beforehand that we were measuring language processing during interaction), four participants reported 'reaction to expectation violations and/or unexpected events'. Two participants reported to the goal question 'finding of a new word'. The remaining participants either stated the known purpose (language processing during interaction) or measuring electrophysiological activity, memory, and/or processing between brain and speech. All participants confirmed that they did not know the confederate before the experiment.

Further, participants were asked to rate on a five-point scale from 'not at all' to 'very' naturalness and pleasantness aspects of the paradigm. We translated these responses to 'not at all' equals 0, 'a bit' equals 1, 'neutral' equals 2, 'somewhat' equals 3, and 'very much' equals 4. On average, the participants rated the communication as somewhat natural ($Mdn$ = 3 ± 0.99), ranging from 0 (n=1) to 4 (n=2). The naturalness of the interaction was rated similarly ($Mdn$ = 3 ± 0.67), ranging from 1 (n=1) to 4 (n=1). The setup was rated as rather pleasant ($Mdn$ = 3 ± 0.77), similar to the pleasantness of the microphones ($Mdn$ = 4 ± 0.79) and room of the experiment ($Mdn$ = 3 ± 0.72). The interaction was rated as somewhat pleasant ($Mdn$ = 3 ± 0.75), ranging from 2 (n=4) to 4 (n=10). Moreover, the interaction partner (i.e., the confederate) was rated as very pleasant ($Mdn$ = 4 ± 0.65), ranging from 2 (n=2) to 4 (n=18), similar to the rating of the pleasantness of her voice ($Mdn$ = 4 ± 0.66) and pleasantness of the experimenter ($Mdn$ = 4 ± 0.56). All participants reported that they were able to make sense of the constructed sentences ($Mdn$ = 3 ± 0.65), ranging from 2 (n=3) to 4 (n=7).

#### GLMM & LMM results

Supplementary table 1

*GLMM results*

| ***Fixed effects*** | Model summary | | | | Model comparison | | |
|---|---|---|---|---|---|---|---|
| | β | SE | t-value | predicted turntime back-transformed | χ2 | df | p-value |
| **Final model** | | | | | | | |
| Intercept | 1.87 | 0.22 | 8.68 | 0.53 | | | |
| Expectancy | -0.79 | 0.17 | -4.65 | 0.92 | 83.88 | 3 | **< .001** |
| Interindividual frequency | 0.98 | 0.15 | 6.66 | 0.35 | 43.11 | 2 | **< .001** |
| Interindividual frequency * Expectancy | -1.10 | 0.17 | -6.60 | 1.30 | 40.98 | 1 | **< .001** |
| **First model** | | | | | | | |
| Intercept | 2.01 | 0.26 | 7.73 | 0.50 | | | |
| Expectancy | -0.77 | 0.17 | -4.56 | 0.81 | | | |
| Intraindividual frequency (1) | -0.13 | 0.16 | -0.82 | 0.53 | | | |
| Intraindividual frequency (2) | -0.17 | 0.16 | -1.11 | 0.54 | | | |
| Interindividual frequency | 1.02 | 0.15 | 6.76 | 0.33 | | | |
| Interindividual frequency * Expectancy | -1.12 | 0.17 | -6.65 | 1.12 | | | |

| ***Random effects*** | Variance | Correlation |
|---|---|---|
| **Final model** | | |
| Wordlength | 0.038 | |
| Participant | 0.431 | |
| Expectancy in participant | 0.222 | -0.091 |
| **First model** | | |
| Wordlength | 0.036 | |
| Participant | 0.432 | |
| Expectancy in participant | 0.222 | -0.091 |

*Note* : GLMM was fitted with a Gamma probability distribution and an inverse link function. Coefficients and SE are NOT backtransformed. The predicted turntime is the backtransformed turntime value in seconds: intercept is the overall mean predicted turntime, mean predicted turntime for expectancy is for moving from expected to unexpected condition (all other factors remaining constant), for intraindividual frequency of naming for moving from less stable to more stable naming, for interindividual frequency of naming is for moving from the less frequent to more frequent (all other factors remaining constant). Significant effects are highlighted in bold.

Supplementary table 2

*LMM results N400*

| ***Fixed effect*** | Model summary | | | Model comparison | | |
|---|---|---|---|---|---|---|
| | β | SE | t-value | χ2 | df | p-value |
| *Anterior midline* | | | | | | |
| Intercept | 1.74 | 0.40 | 4.31 | | | |
| Expectancy | -0.51 | 0.28 | -1.85 | 3.42 | 1 | 0.064 |
| *Central midline* | | | | | | |
| Intercept | 2.37 | 0.37 | 6.37 | | | |
| Expectancy | -1.52 | 0.32 | -4.75 | 16.53 | 1 | **< .001** |
| *Posterior midline* | | | | | | |
| Intercept | 1.44 | 0.26 | 5.50 | | | |
| Expectancy | -1.45 | 0.32 | -4.55 | 15.48 | 1 | **< .001** |
| *Left anterior quadrant* | | | | | | |
| Intercept | 1.12 | 0.31 | 3.62 | | | |
| Expectancy | -0.29 | 0.25 | -1.15 | 1.33 | 1 | 0.249 |
| *Left posterior quadrant* | | | | | | |
| Intercept | 1.72 | 0.28 | 6.11 | | | |
| Expectancy | -1.41 | 0.24 | -5.96 | 35.14 | 1 | **< .001** |
| *Right anterior quadrant* | | | | | | |
| Intercept | 1.16 | 0.30 | 3.88 | | | |
| Expectancy | -0.46 | 0.23 | -1.99 | 3.94 | 1 | **0.047** |
| *Right posterior quadrant* | | | | | | |
| Intercept | 1.19 | 0.24 | 5.01 | | | |
| Expectancy | -1.30 | 0.22 | -5.86 | 34.02 | 1 | **< .001** |

| ***Random effects*** | Variance | Correlation |
|---|---|---|
| *Anterior midline* | | |
| Participant | 0.00 | |
| Expectancy in participant | 0.00 | 1 |
| *Central midline* | | |
| Participant | 2.41 | |
| Expectancy in participant | 0.52 | -0.04 |
| *Posterior midline* | | |
| Participant | 0.92 | |
| Expectancy in participant | 0.89 | -0.23 |
| *Left anterior quadrant* | | |
| Participant | 1.71 | |
| Expectancy in participant | 0.27 | -0.14 |
| *Left posterior quadrant* | | |
| Participant | 1.28 | |
| *(failed to converge with slope)* | | |
| *Right anterior quadrant* | | |
| Participant | 1.56 | |
| Expectancy in participant | 0.022 | 1 |
| *Right posterior quadrant* | | |
| Participant | 0.81 | |
| *(failed to converge with slope)* | | |

*Note:* LMM was fitted with expectancy (expected vs. unexpected) as fixed factor and random slope expectancy in participant and random intercept participant. Significant effects are highlighted in bold. Model for left and right posterior quadrant with random slope congruency in participant failed to converge, therefore, simpler model with participant as random intercept is reported.

Supplementary table 3

*LMM results P600*

| ***Fixed effect*** | Model summary | | | Model comparison | | |
|---|---|---|---|---|---|---|
| | β | SE | t-value | χ2 | df | p-value |
| *Anterior midline* | | | | | | |
| Intercept | -0.21 | 0.31 | -0.66 | | | |
| Expectancy | -0.19 | 0.27 | -0.68 | 0.46 | 1 | 0.497 |
| *Central midline* | | | | | | |
| Intercept | 0.53 | 0.30 | 1.77 | | | |
| Expectancy | 0.45 | 0.28 | 1.62 | 2.61 | 1 | 0.106 |
| *Posterior midline* | | | | | | |
| Intercept | 0.97 | 0.26 | 3.78 | | | |
| Expectancy | 0.85 | 0.27 | 3.18 | 8.66 | 1 | **0.003** |
| *Left anterior quadrant* | | | | | | |
| Intercept | 0.03 | 0.23 | 0.14 | | | |
| Expectancy | -0.31 | 0.25 | -1.26 | 1.61 | 1 | 0.205 |
| *Left posterior quadrant* | | | | | | |
| Intercept | 0.96 | 0.23 | 4.12 | | | |
| Expectancy | 0.63 | 0.24 | 2.60 | 6.40 | 1 | **0.011** |
| *Right anterior quadrant* | | | | | | |
| Intercept | -0.08 | 0.26 | -0.31 | | | |
| Expectancy | -0.45 | 0.22 | -2.00 | 3.96 | 1 | **0.046** |
| *Right posterior quadrant* | | | | | | |
| Intercept | 0.65 | 0.24 | 2.71 | | | |
| Expectancy | 0.60 | 0.21 | 2.89 | 8.34 | 1 | **0.004** |

| ***Random effects*** | Variance | Correlation |
|---|---|---|
| *Anterior midline* | | |
| Participant | 1.58 | |
| Expectancy in participant | 0.07 | -1.00 |
| *Central midline* | | |
| Participant | 1.34 | |
| Expectancy in participant | 0.03 | 1.00 |
| *Posterior midline* | | |
| Participant | 0.94 | |
| Expectancy in participant | 0.29 | 0.26 |
| *Left anterior quadrant* | | |
| Participant | 0.69 | |
| Expectancy in participant | 0.30 | -0.46 |
| *Left posterior quadrant* | | |
| Participant | 0.74 | |
| Expectancy in participant | 0.20 | 1.00 |
| *Right anterior quadrant* | | |
| Participant | 1.02 | |
| Expectancy in participant | 0.02 | -1.00 |
| *Right posterior quadrant* | | |
| Participant | 0.89 | |
| *(failed to converge with slope)* | | |

*Note:* LMM was fitted with expectancy (expected vs. unexpected) as fixed factor and random slope expectancy in participant and random intercept participant. Significant effects are highlighted in bold. Model for right posterior quadrant failed to converge, therefore, simpler model with only random intercept participant is reported.

Supplementary table 4

*GLMM results*

Turntime explained by N400 EEG activity

| ***Fixed effects*** | Model summary | | | | Model comparison | | |
|---|---|---|---|---|---|---|---|
| | β | SE | t-value | predicted turntime back-transformed | χ2 | df | p-value |
| *Anterior midline* | | | | | | | |
| Intercept | 2.36 | 0.21 | 11.02 | 0.424 | | | |
| N400 EEG activity | 0.01 | 0.01 | 1.86 | 0.421 | 43.13 | 3 | **< .001** |
| Expectancy | -1.44 | 0.14 | -10.33 | 1.088 | 42.94 | 2 | **< .001** |
| N400 EEG activity * expectancy | -0.02 | 0.01 | -1.87 | 0.427 | 3.40 | 1 | 0.065 |
| *Central midline* | | | | | | | |
| Intercept | 2.35 | 0.21 | 11.04 | 0.426 | | | |
| N400 EEG activity | 0.02 | 0.01 | 2.97 | 0.422 | 53.05 | 3 | **< .001** |
| Expectancy | -1.43 | 0.14 | -10.29 | 1.086 | 42.03 | 2 | **< .001** |
| N400 EEG activity * expectancy | -0.01 | 0.01 | -1.67 | 0.429 | 2.71 | 1 | 0.100 |
| *Posterior midline* | | | | | | | |
| Intercept | 2.37 | 0.21 | 11.23 | 0.423 | | | |
| N400 EEG activity | 0.03 | 0.01 | 3.18 | 0.418 | 68.36 | 3 | **< .001** |
| Expectancy | -1.43 | 0.14 | -10.37 | 1.067 | 39.86 | 2 | **< .001** |
| N400 EEG activity * expectancy | -0.01 | 0.01 | -0.93 | 0.424 | 0.84 | 1 | 0.359 |
| *Left anterior quadrant* | | | | | | | |
| Intercept | 2.38 | 0.21 | 11.10 | 0.420 | | | |
| N400 EEG activity | 0.00 | 0.01 | -0.02 | 0.420 | 40.95 | 3 | **< .001** |
| Expectancy | -1.46 | 0.14 | -10.45 | 1.087 | 39.88 | 2 | **< .001** |
| N400 EEG activity * expectancy | -0.01 | 0.01 | -0.50 | 0.421 | 0.25 | 1 | 0.617 |
| *Left posterior quadrant* | | | | | | | |
| Intercept | 2.36 | 0.21 | 11.17 | 0.423 | | | |
| N400 EEG activity | 0.02 | 0.01 | 2.49 | 0.419 | 56.09 | 3 | **< .001** |
| Expectancy | -1.43 | 0.14 | -10.34 | 1.076 | 39.80 | 2 | **< .001** |
| N400 EEG activity * expectancy | -0.01 | 0.01 | -0.79 | 0.425 | 0.62 | 1 | 0.433 |
| *Right anterior quadrant* | | | | | | | |
| Intercept | 2.37 | 0.21 | 11.07 | 0.422 | | | |
| N400 EEG activity | 0.01 | 0.01 | 1.27 | 0.420 | 41.26 | 3 | **< .001** |
| Expectancy | -1.45 | 0.14 | -10.42 | 1.089 | 41.19 | 2 | **< .001** |
| N400 EEG activity * expectancy | -0.01 | 0.01 | -1.30 | 0.424 | 1.65 | 1 | 0.199 |
| *Right posterior quadrant* | | | | | | | |
| Intercept | 2.36 | 0.21 | 11.26 | 0.424 | | | |
| N400 EEG activity | 0.04 | 0.01 | 3.95 | 0.417 | 73.34 | 3 | **< .001** |
| Expectancy | -1.43 | 0.14 | -10.43 | 1.072 | 41.67 | 2 | **< .001** |
| N400 EEG activity * expectancy | -0.02 | 0.01 | -1.66 | 0.427 | 2.68 | 1 | 0.101 |

| ***Random effects*** | Variance | Correlation |
|---|---|---|
| *Anterior midline* | | |
| Participant | 0.48 | |
| Wordlength | 0.09 | |
| Expectancy in participant | 0.24 | -0.90 |
| *Central midline* | | |
| Participant | 0.48 | |
| Wordlength | 0.08 | |
| Expectancy in participant | 0.24 | -0.91 |
| *Posterior midline* | | |
| Participant | 0.47 | |
| Wordlength | 0.08 | |
| Expectancy in participant | 0.24 | -0.90 |
| *Left anterior quadrant* | | |
| Participant | 0.48 | |
| Wordlength | 0.09 | |
| Expectancy in participant | 0.24 | -0.90 |
| *Left posterior quadrant* | | |
| Participant | 0.48 | |
| Wordlength | 0.08 | |
| Expectancy in participant | 0.24 | -0.90 |
| *Right anterior quadrant* | | |
| Participant | 0.48 | |
| Wordlength | 0.09 | |
| Expectancy in participant | 0.24 | -0.90 |
| *Right posterior quadrant* | | |
| Participant | 0.46 | |
| Wordlength | 0.08 | |
| Expectancy in participant | 0.23 | -0.90 |

*Note* : GLMM was fitted with a Gamma probability distribution and an inverse link function. Coefficients and SE are NOT backtransformed. The predicted turntime is the backtransformed turntime value in seconds. Significant effects are highlighted in bold.

**Speech onset times relative to sentence start**

Speech onset times in a sentence over all trials are shown in supplementary figure 2. We can see that the onset of the first word is the sharpest in time relative to sentence onset, meaning that the confederate had similar reaction times to utter the first word. The histogram displays the overall distribution of the word onsets within the sentence, where there is a division between expected and unexpected fourth words uttered by the participant. We see that fourth words following an unexpected article are uttered later in time and have a right skewed distribution as compared to fourth words following an expected article.

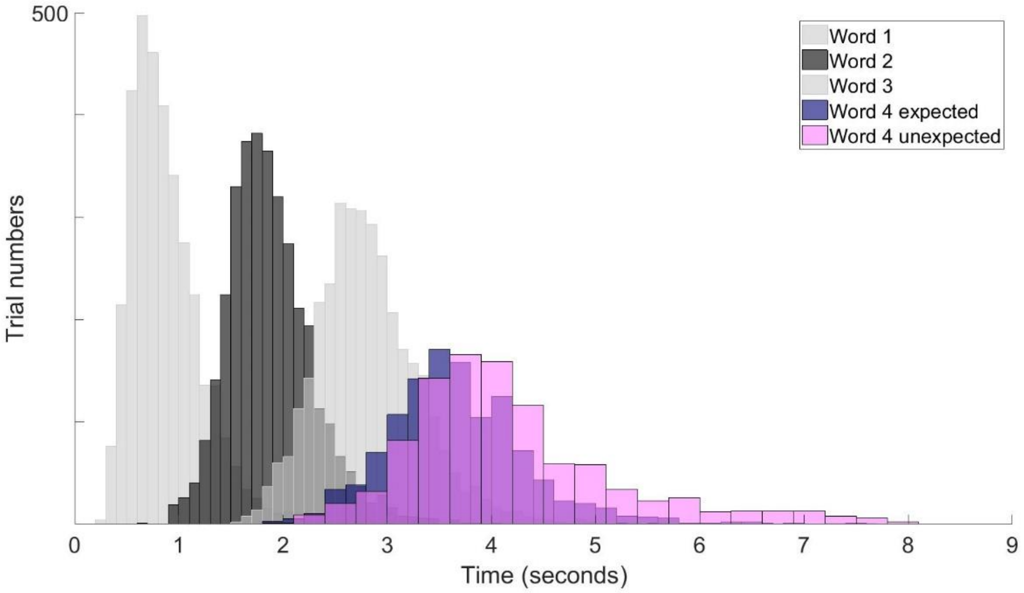


**Supplementary figure 2**: Speech onset times for word 1 to word 4 over the period of a sentence. Zero point is the onset of the sentence. Values below 0 or above 8 seconds were not included. Word 1 and 3 (light grey) were uttered by the confederate. Word 2 (dark grey) and word 4 (expected in blue and unexpected in magenta) were uttered by the participant.

**Grand average ERPs to confederate-produced words**

The grand average ERPs over all conditions for the first name and the article (word 1 and word 3, see supplementary fig. 3) show a typical response to auditory speech stimuli: an increased amplitude in the N100 time window and a topographic distribution typical for the N100 (May & Tiitinen, 2010), followed by a short positivity resembling a P200 response and topography (Goregliad Fjaellingsdal et al., 2016). An N400 typical response is apparent for both heard words but at different latencies, earlier for word 3 compared to word 1. We relate this latency difference to the significantly different speech lengths of word 1 and word 3, a first name vs. an article ($t(46) = 27.90$, $p < .001$; see supplementary fig.3). The topography for the N400 for word 3 is shown earlier due to the average length of the confederate produced word (~400 ms). The focal distribution is similar between the different time points (T3) for word 1 and word 3, though with different relative activity (negative for word 1 and positive for word 3).

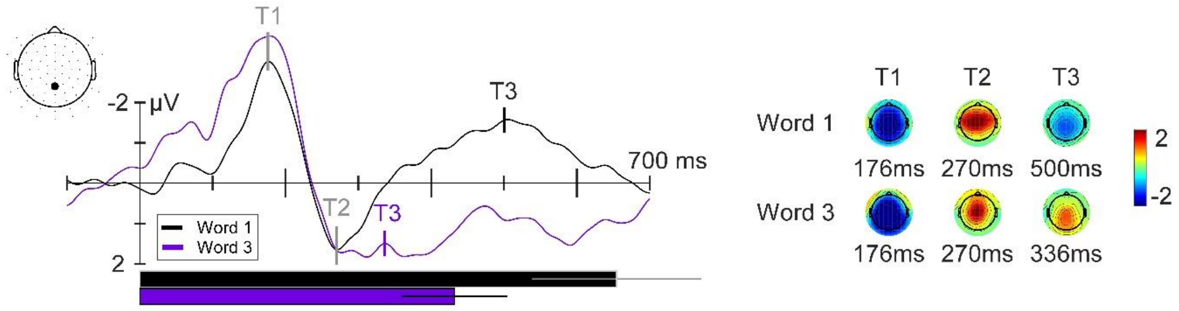


**Supplementary figure 3**: Grand average ERP in reaction to the confederate-produced words, word 1 (black) and word 3 (purple), for all conditions and participants at channel 05 (see highlighted electrode on layout map). Peak topographies are shown for the N100 at 176 ms (T1), for the P200 at 270 ms (T2), and for the N400 at 500 ms for word 1 (T3 in black) and 336 ms for word 3 (T3 in purple). Statistical analysis on the critical trials of word 3 (split for expectancy conditions) was run for the N400 between 250 and 450 ms, so T3 for word 3 is set here approximately at the middle of this time span. Peak topographies' time points are depicted with a vertical line in the ERP plot. Zero point is the onset of the words. Mean word duration (of confederate) plus standard deviation is shown with the same time scale below the ERP.

**Global map dissimilarity**

To assess topographical differences, global map dissimilarity (Michel, Koenig, Brandeis, Gianotti, & Wackermann, 2009, p. 45) between expected and unexpected conditions was calculated over time for the ERP of word 3 (see supplementary figure 4). We found high similarities (shown in red colour) for the N100 time window until around 220 ms after word onset. Notably, dissimilarity between the two expectancy conditions reaches a peak between 220 to 450 ms. Whereas the topography for the expected condition shows a stable positive deflection over this time period, a negative and positive deflection are visible for the unexpected condition (see supplementary video). Dissimilarity is apparent again within the P600 time window from 500 to 700 ms, where we see different topographical maps between expectancy conditions.

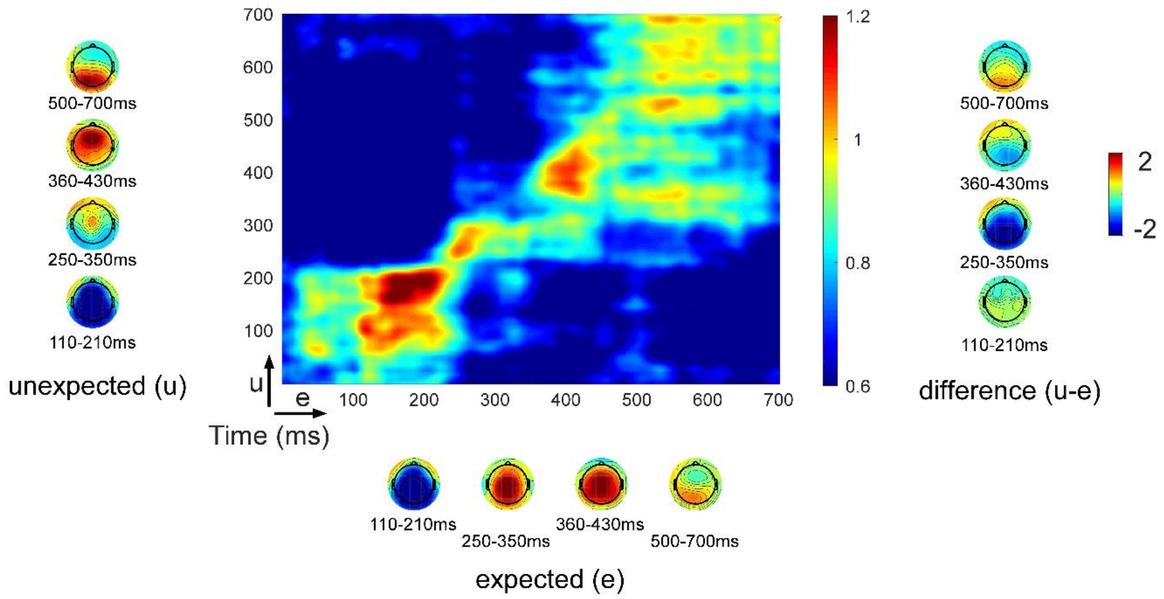


**Supplementary figure 4**: Global map dissimilarity (GMD) power calculated over time between expected and unexpected conditions (ERP of word 3). Higher values indicating here higher similarity of topographic distributions. Grand average topographies for specific time ranges are shown for each condition next to its axis (x-axis = expected; y-axis = unexpected) and their difference topographies for the same time windows on the right side of the figure. Topography scale applies to all shown topographies.

#### Supplementary video

**Supplementary video**: Video of ERP activity for word 3: expected condition (upper row), unexpected condition (middle row), and the difference unexpected minus expected (lower row). Left column shows the topographic activity for the time period from onset of the word at 0 ms to 700 ms after onset. Right column shows the ERP plot for each channel for expected and unexpected conditions. The ERP plot of the difference condition shows the activity at channel 05 (highlighted on the topography) for expected (blue), unexpected (magenta), and their difference (black).
